# Proteomic Analyses of Morphological Variants of *Borrelia burgdorferi* Shed New Light on Persistence Mechanisms: Implications for Pathogenesis, Diagnosis and Treatment

**DOI:** 10.1101/501080

**Authors:** Jie Feng, Ying Zhang

## Abstract

*Borrelia burgdorferi* causes Lyme disease, which is the most common vector borne disease in the United States and Europe. Although 2-4 week antibiotic treatment for Lyme disease is effective in the majority of cases, about 10–20% patients suffer from prolonged post-treatment Lyme disease syndrome (PTLDS). While the mechanisms of PTLDS are unclear, persisting organisms not killed by current Lyme antibiotics has been suggested as a possible explanation. *B. burgdorferi* can spontaneously develop different morphological variant forms under stress or in stationary phase with increased persistence to antibiotics. To shed light on the possible mechanisms by which these variant forms develop persistence, here, we isolated three *B. burgdorferi* forms, log phase spirochetal form, stationary phase planktonic form, and stationary phase aggregated biofilm-like microcolony form. We showed that the two separated stationary phase forms especially microcolony form have more persistence to antibiotics than the log phase spirochetal form. Then, we performed mass spectrometry (MS/MS) analysis to determine the proteomic profiles of the three different forms to reveal the mechanisms of persistence in *B. burgdorferi*. We identified 1023 proteins in the three *B. burgdorferi* forms, with 642 proteins (63%) differentially expressed. Compared with the log phase spirochetal form of *B. burgdorferi*, a total of 143 proteins were upregulated in both stationary phase planktonic form and microcolony form. Among these common upregulated proteins, 90 proteins had predicted functions and were mapped to different pathways involved in infection and virulence, DNA repair, heat shock, transport, sporulation, cell envelope and metabolism, many of which are consistent with persister mechanisms in other bacteria. A particularly interesting observation is that infection and virulence related proteins are highly up-regulated in stationary phase planktonic form and microcolony form compared with log phase spirochetal form. These findings shed new light on the mechanisms of *B. burgdorferi* persistence and offer novel targets for developing more effective diagnostics, vaccines and treatments.

## Introduction

Lyme disease (LD) is the most prevalent tick-borne illness and an important emerging zoonosis in the United States with an estimated 300,000 cases per year [1]. The causative agent of Lyme disease is pathogenic Borrelia species including *B. burgdorferi, B. afzelii, B. garinii*. *B. burgdorferi* is the predominant cause of human LD in North America. In humans, Lyme disease may cause a local erythema migrans rash at the site of the tick bite and then readily disseminates through the bloodstream to other tissues, setting up an infection that can last for months to years. Patients with Lyme disease are routinely treated with doxycycline, amoxicillin or cefuroxime, which effectively hastens the resolution in most cases. However, about 10%-20% patients continue to suffer lingering symptoms of fatigue, pain or joint and muscle aches, and neurocognitive manifestations that last 6 months or more despite treatment, a condition called “post-treatment Lyme disease syndrome” (PTLDS) [2]. While the cause of PTLDS is complex and remains to be determined, one of the possible explanations is persistent *B. burgdorferi* infection due to persisters not killed by the current antibiotic treatment, which evade host immune clearance and drive immunological responses continually as shown in various animal models [3–6]. A recent study in humans demonstrated the recovery of *B. burgdorferi* DNA by xenodiagnoses in patients despite antibiotic treatment [7]. Findings indicate that current Lyme disease treatment may not sufficiently eliminate *B. burgdorferi* persisters or that the immune system fails to clear persisting organisms or bacterial debris, which may be the underlying cause for those who suffer from unresolved Lyme disease symptoms. In contrast to other bacterial pathogens that cause persistent infections such as *M. tuberculosis* and *E. coli,* an unusual feature of the in vivo persistence of *B. burgdorferi* is the lack of culturability of the persisting organisms despite the demonstration of its DNA and even increased DNA content by PCR or by xenodiagnosis [3, 4, 8].

In addition to the above in vivo persistence, *B. burgdorferi* has recently been shown to develop persisters in vitro in cultures as shown by tolerance to current Lyme antibiotics doxycycline, amoxicillin and cefuroxime [9–11]. In addition, *B. burgdorferi* can develop morphological variants including spirochetal form, round body form, or cystic form, and aggregated biofilm-like microcolonies as the culture grows from log phase to stationary phase or under stress conditions in vitro [9, 12]. The variant forms such as cystic and round body forms have been found in vivo in brain tissues of Lyme borreliosis patients [13] but their role in persistent form of the disease is controversial [14]. Frontline drugs such as doxycycline and amoxicillin could kill the replicating spirochetal form of *B. burgdorferi* quite effectively, but they exhibit poor activity in killing non-replicating persister forms [9–11]. We showed that stationary phase cultures contain different morphological variants including planktonic spirochetal form, round body form and aggregated microcolony form [9, 12], which have varying levels of persistence or can be considered different types of persisters in comparison to the log phase culture which mainly consists of growing spirochetal form with no or few persisters [12]. Therefore, it is critical to understand the mechanism of the decreased drug susceptibility or persistence in the morphological variants of *B. burgdorferi*, which will be important for developing drugs targeting such forms for effective treatments. Previously, we have analyzed the transcriptome of *B. burgdorferi* persisters by RNA-seq and identified a range of genes and pathways involved in persistence to doxycycline and amoxicillin [15]. However, little is known about the proteomic profiles of the different forms or persisters of *B. burgdorferi*. As part of our ongoing study to understand the molecular basis of persistence in *B. burgdorferi,* we isolated three different forms of *B. burgdorferi*, including growing log phase spirochetes (as a control), non-growing planktonic form and aggregated biofilm-like microcolony form and subjected them to proteomic analysis. To quantitatively compare the changes in protein expression of three different forms of *B. burgdorferi*, in this study, we used tandem mass spectrometry (MS/MS) and NSAF (normalized spectral abundance factor) method to analyze protein abundance of samples of different *B. burgdorferi* forms. Our findings not only shed new light on *B. burgdorferi* persistence mechanisms in this intriguing and versatile organism but also offer new targets for intervention. To our knowledge, this is the first report of proteomics-based analyses of different morphological variants of *B. burgdorferi*.

## Materials and Methods

### Strain, media, and culture techniques

The low passaged (less than 5 passages) *Borrelia burgdorferi* strain B31 5A19 was kindly provided by Dr. Monica Embers [11]. *B. burgdorferi* strains were grown in BSK-H medium (HiMedia Laboratories Pvt. Ltd.) with 6% rabbit serum (Sigma-Aldrich, St. Louis, MO, USA). All culture medium was filter-sterilized by 0.2 μm filter. Cultures were incubated in microaerophilic incubator (33°C, 5% CO_2_) without antibiotics.

### Antibiotics and drug susceptibility assay

Antibiotics including doxycycline, ceftriaxone and cefuroxime were purchased from Sigma-Aldrich (St. Louis, MO, USA) and dissolved in water at appropriate concentrations to form stock solutions. All antibiotic stocks were filter-sterilized using a 0.2 μm filter. The residual viability of *B. burgdorferi* cells treated with antibiotics or drug combinations were evaluated using the SYBR Green I/PI assay combined with fluorescence microscopy as described previously [16,22]. Briefly, the ratio of live and dead cells was calculated by the ratio of green/red fluorescence and the regression equation and regression curve with least-square fitting analysis.

### Separation and preparation of microcolony form and planktonic form from stationary phase culture

After incubation for 10 days, 1 ml stationary phase *B. burgdorferi* culture (∼10^7^ spirochetes/mL) was centrifuged at 800 × g for 10 min, and the supernatant was transferred to a new tube as stationary phase spirochetal form. The bottom 50 μl microcolony rich culture was resuspended and centrifuged at 800 × g 3 times to remove the planktonic spirochetes. The stationary phase spirochetal form and microcolony form were checked with fluorescence microscope to ensure their morphologies before being used for the infection and mass spectrum analysis (see below).

### Preparation of protein samples

*B. burgdorferi* 3-5-day old log phase cells, 10-day old stationary phase spirochetal cells and stationary phase microcolony cells were collected by centrifugation at 9000 × g for 10 min. The cells were washed with 4°C phosphate buffered saline (PBS) 5 times to remove serum proteins in the BSK medium. Then the cells were resuspended in 0.5 ml 2% SDS containing 1 mM EDTA and 1 mM PMSF, and sonicated. Debris were removed by centrifugation. Small portion of the supernatant was diluted and measured for total protein concentration. Aliquoted samples were stored at −80°C or submitted on dry ice for proteomic studies and other experiments.

### Nanospray LC/MS/MS and database search

The protein samples were concentrated and separated by SDS-PAGE. The gel pieces were subjected to in-gel trypsin digestion and LC/MS/MS analysis. The LC/MS/MS analysis of samples was carried out using a Thermo Fisher Scientific Q-Exactive hybrid Quadrupole-Orbitrap Mass Spectrometer and a Thermo Dionex UltiMate 3000 RSLCnano System. Eight tryptic peptide mixtures from eight gel-pieces from each sample were analyzed by LC/MS/MS. For each LC/MS/MS run the tryptic peptide mixture was loaded onto a peptide trap cartridge at a flow rate of 5 μL/min. The trapped peptides were eluted onto a reversed-phase PicoFrit column (New Objective, Woburn, MA) using a linear gradient of acetonitrile (3-36%) in 0.1% formic acid. The elution duration was 60 min at a flow rate of 0.3 μl/min. Eluted peptides from the PicoFrit column were ionized and sprayed into the mass spectrometer, using a Nanospray Flex Ion Source ES071 (Thermo Fisher Scientific) under the following settings: spray voltage, 1.6 kV, Capillary temperature, 250°C. For protein identification two raw MS files from two LC/MS/MS runs for each sample were analyzed using the Thermo Proteome Discoverer 1.4.1 platform (Thermo Scientific, Bremen, Germany) for peptide identification and protein assembly. Database search against NCBI public *Borrelia burgdorferi* B31 protein database (4679 entries) obtained from NCBI website was performed based on the SEQUEST and percolator algorithms through the Proteome Discoverer 1.4.1 platform. Carbamidomethylation of cysteines was set as a fixed modification, and Oxidation, Deamidation Q/N-deamidated (+0.98402 Da) were set as dynamic modifications. The minimum peptide length was specified to be five amino acids. The precursor mass tolerance was set to 15 ppm, whereas fragment mass tolerance was set to 0.05 Da. The maximum false peptide discovery rate was specified as 0.05. The resulting Proteome Discoverer Report contains all assembled proteins with peptide sequences and matched spectrum counts. The estimation of relative abundance of proteins was based on peptide spectrum match counts (PSM).

### Protein quantification

Protein quantification used the normalized spectral abundance factors (NSAFs) method [16, 17] to calculate the protein relative abundance. In order to quantitatively describe the relative abundance, the ppm (part per million) was chosen as the unit and the 1,000,000 ppm value was assigned to each proteome profile. A ppm value in the range of 0 to 1,000,000 ppm for each identified protein in each proteome profile was calculated based on its normalized NSAF.

### Data analysis and metabolic pathway construction

All detected proteins were imported into a local MySQL database by custom-made script for analysis. The KEGG database [18] was used to map the detected proteins to metabolism pathways with custom-made MySQL script. For pathways in which some functions were missing, related proteins were searched against nr database (containing all non-redundant sequences from GenBank CDS translations, PDB, Swiss-Prot, PIR, and PRF) using BLAST program [19] to identify additional functions.

## Results

### Separation of aggregated microcolony form from planktonic form in stationary phase culture

Previous studies demonstrated that *B. burgdorferi* develops multiple morphological forms including spirochetal form, round bodies (cysts), and aggregated microcolony form [9, 13]. Log phase *B. burgdorferi* culture is mainly in spirochetal form, but stationary phase culture is dominated by coccoid or round-body forms and aggregated micro-colony forms [9]. We showed that the current clinically used antibiotics used for treating Lyme disease had high activity against log phase *B. burgdorferi* but had very limited activity against the stationary phase cells [9]. We also demonstrated that aggregated microcolony forms are more tolerant to antibiotics than planktonic spirochetal and round body forms in the stationary phase culture [12]. To identify proteins that are preferentially expressed in the variant forms to shed new light on persistence mechanisms, here we first separated the aggregated microcolony form from the planktonic form (spirochetal and round body) in stationary phase *B. burgdorferi* culture according to their different densities using low speed centrifugation. After multiple differential centrifugation separation, microcolony form and planktonic form were separated effectively (Figure 1B and C). Consistent with our previous study [12], here we found that the 5 day old log phase cells were mostly in spirochetal form (Figure 1A), while the 10-day old stationary phase culture contained, in addition to aggregated biofilm-like microcolonies (Fig. 1B), planktonic forms made up of not only spirochetes but also many round body cells (Figure 1C).

**Figure 1.**
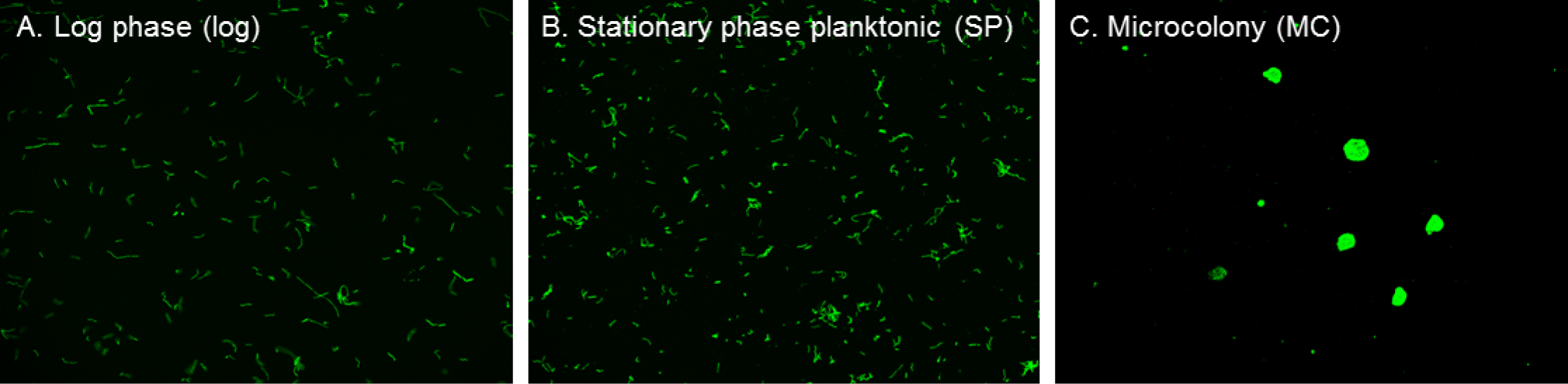
Representative images of a 3-day old log phase spirochete form (A), 10-day old stationary phase, 10-day stationary phase planktonic form (B) and aggregated microcolony form (C) of *B. burgdorferi.* Cells were stained with SYBR Green I/PI viability assay and observed using fluorescence microscopy at 200 X magnification.

### Overall features of proteome profiles of different variant forms of *B. burgdorferi* revealed by LC/MS/MS analysis

Total proteins from microcolony form (MC) (Fig. 1B), stationary phase planktonic form (SP) (Fig. 1C) and log phase spirochetal form (LOG) (Fig. 1A) were analyzed by LC/MS/MS. PSM counts are proportional to the protein amount analyzed in the three separate analyses. A total of 1023 proteins were identified in the three analyses, with 642 proteins (63%) differentially expressed among three samples (>= 2-fold change, CV>75%, >3 PSMs). To validate the differentially expressed proteins, 62 house-keeping proteins such as 30S and 50S ribosomal proteins, DNA polymerases, DNA-directed RNA polymerases, and a few other proteins were used as the internal control. The majority of selected house-keeping proteins were at the same level or the change in terms of the CV was less than 50% among the three samples. There are 55 ribosomal proteins (22 30S proteins and 33 50S proteins) in the *B. burgdorferi* B31 protein database, and 48 of them were identified in all three samples, suggesting the coverage for the ribosomal proteins in our samples is quite good at 87%.

### Comparison of proteome profiles of stationary phase planktonic form and log phase *B. burgdorferi*

Although the stationary phase planktonic form was similar to log phase cells in morphology (Figure 1), their proteome profiles were very different. We found 188 proteins were up-regulated in the stationary phase planktonic form compared to the log phase *B. burgdorferi* (Table S1). Among them, the functions of 87 proteins are predicted (Table S1). We found that 15 proteins are involved in the metabolism of nucleotide, carbohydrate, amino acids, etc. (Figure 2). Additionally, 16 infection and virulence proteins are up-regulated in the stationary phase planktonic form compared to the log phase *B. burgdorferi* cells, which include 10 Bdr proteins (BdrAEFHMOPRVW), two decorin-binding proteins (DbpAB), a ChpAI protein (BB_A07), a complement regulator-acquiring surface protein (CRASP, BB_A68) and a virulence associated lipoprotein BB_I29. We also noticed that the expression of DbpA and DbpB protein was significantly upregulated (4.3 and 7.8 fold, respectively). In addition, we found 8 DNA replication and repair proteins and 7 transporter proteins were upregulated in the stationary phase planktonic form. Furthermore, this study identified 5 molecular chaperones or heat shock proteins including two Clp protease proteins (ClpC and ClpP). In addition, 4 motility and chemotaxis proteins, 17 ribosome proteins, 3 transcriptional regulators, two rRNA modification factors and 9 lipoproteins and membrane proteins were found upregulated in the stationary phase planktonic form (Table S1).

**Figure 2.**
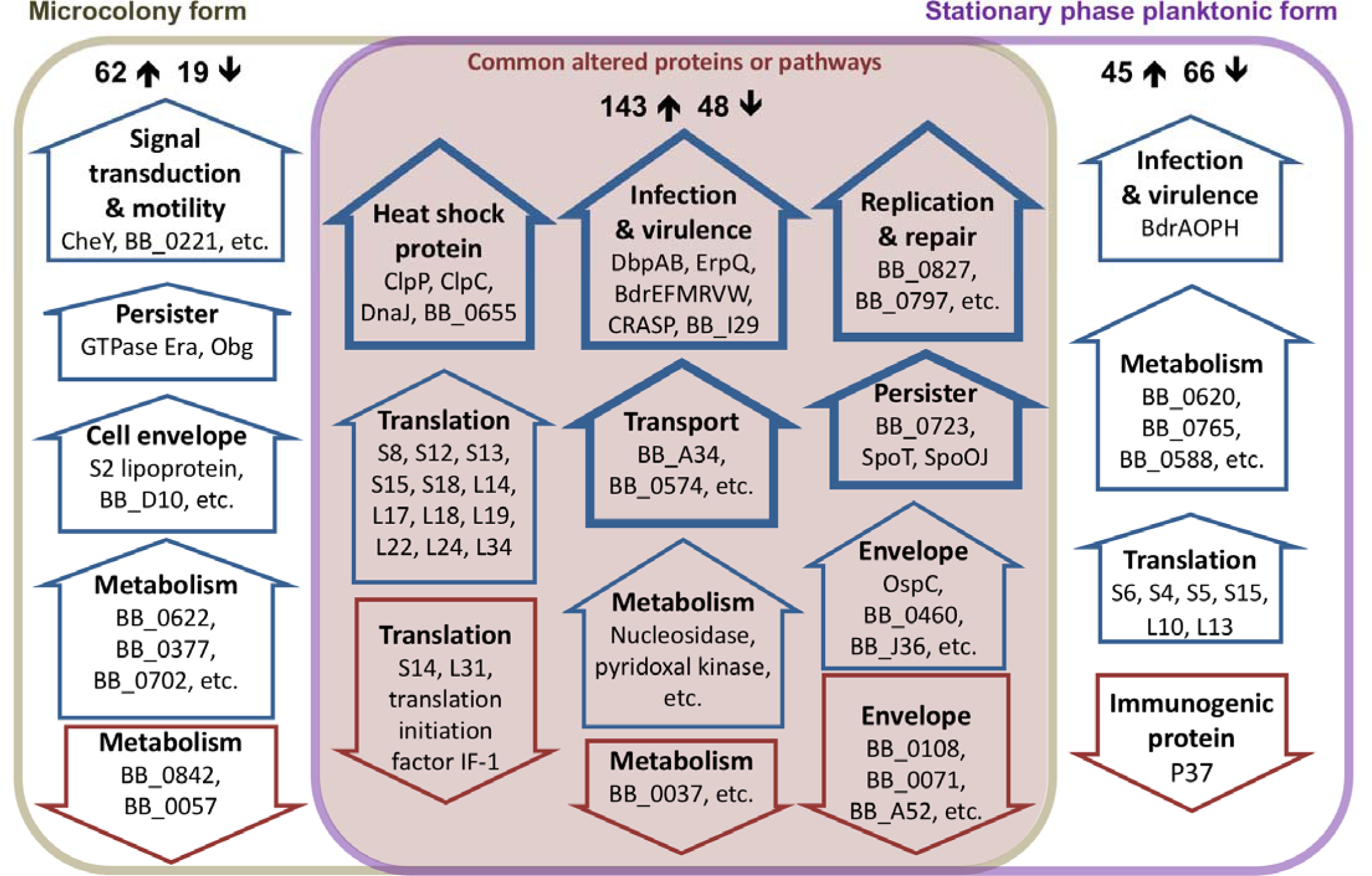
Overlap between the dominating regulated pathways in stationary phase planktonic and microcolony forms compared with log phase *B. burgdorferi*. The blue upward or red downward arrows indicate the up- or down-regulated pathways, respectively. The preferentially active pathways that are identified in both drug-tolerant forms are highlighted in the center pink shaded overlap box. The pathways which are also identified from our previous RNA-seq study on *B. burgdorferi* persisters [15] are highlighted in bold frame. The numbers with arrow show the number of up-regulated or down-regulated proteins in their respective portion.

On the other hand, we found 114 proteins were down-regulated in stationary phase planktonic form compared with log phase culture (Table S1). Among them, 59 proteins have predicted function (Table S1) and belong to several pathways (Figure 2). We found 15 down-regulated proteins are involved in multiple metabolism pathways. Furthermore, many cell motility proteins including 10 flagellar proteins and 3 chemotaxis proteins are dramatically down-regulated in the stationary phase planktonic form; among them 7 proteins (BB_0415, BB_0040, BB_0183, BB_0276, BB_0775, BB_0221, BB_0285) were not detected in the stationary phase planktonic form. Other down-regulated proteins mainly include 3 ribosomal proteins, 3 transporters (BB_B23, BB_B06, BB_0836), two immunogenic proteins (BB_K50, BB_K48), two kinases (BB_0015, BB_0791), DNA methyltransferase (BB_Q67), DNA glycosylase (BB_0053), protein translocase SecD (BB_0652), etc. Interestingly, we did not find Clp protease subunit A (BB_0369) in the stationary phase planktonic form, although the ClpC and ClpP proteins are upregulated.

### Comparison of proteome profiles of microcolony form and log phase *B. burgdorferi*

Microcolony is the most drug-tolerant or persistent form of *B. burgdorferi* [12]. Here we found a total of 205 proteins that were up-regulated (>2-fold change, CV>75%, >3 PSMs) in the microcolony form compared to the log phase *B. burgdorferi*. Among them, 87 proteins with predicted function and pathway are shown in Table S2. We found 19 proteins are involved in various metabolic pathways; and 15 proteins including 13 ribosomal proteins involved in translation (Table 2). Like stationary phase planktonic form, we found 12 up-regulated proteins in the MC form are involved in infection and virulence, which include ChpAI (BB_A07), CRASP (BB_A68), DbpAB, BdrEFMRVW, protein ErpQ and virulence associated lipoprotein BB_I29. In the other up-regulated proteins, we found 10 of them are transporters, 9 proteins related to signal transduction and motility, and 7 proteins involved in DNA replication and repair. Additionally, 12 membrane or lipoproteins are up-regulated in the microcolony form compared with the log phase *B. burgdorferi* cells. Moreover, we also found four molecular chaperone proteins including BB_0655, DnaJ, ClpP and ClpC up-regulated in the microcolony form (Table S2). Besides, we also identified some remarkable persister proteins are up-regulated in the microcolony form, such as BB_0723 adenylyl cyclase [20], GTPase Era [21, 22], GTPase Obg [23], SpoT (guanosine-3’;5’-bis(diphosphate) 3’-pyrophosphohydrolase) [24–26] and sporulation protein spoOJ (Figure 2, Table S2).

Meanwhile, we identified 67 proteins that were down-regulated in the microcolony form compared with log phase culture (Table S2). In the 29 known proteins, we found 7 signal transduction and motility proteins including 5 flagellar proteins. In addition, another 7 down-regulated proteins were found in multiple metabolism pathways. Other down-regulated pathways include RNA modification (5 proteins), translation (4 proteins), DNA replication and repair (2 proteins), one RNA polymerase sigma factor and one chitobiose transporter protein ChbB.

**Table 1.**
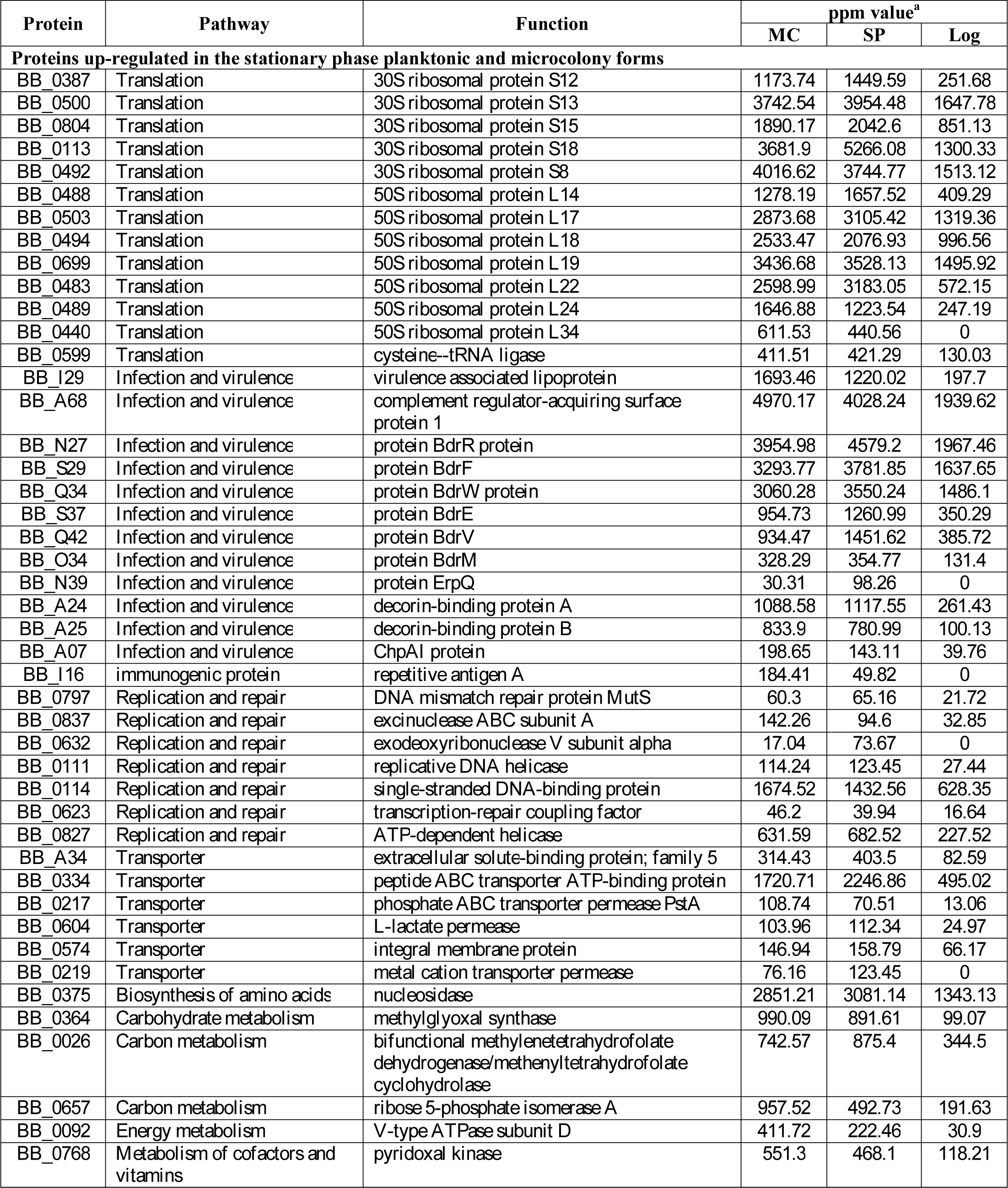
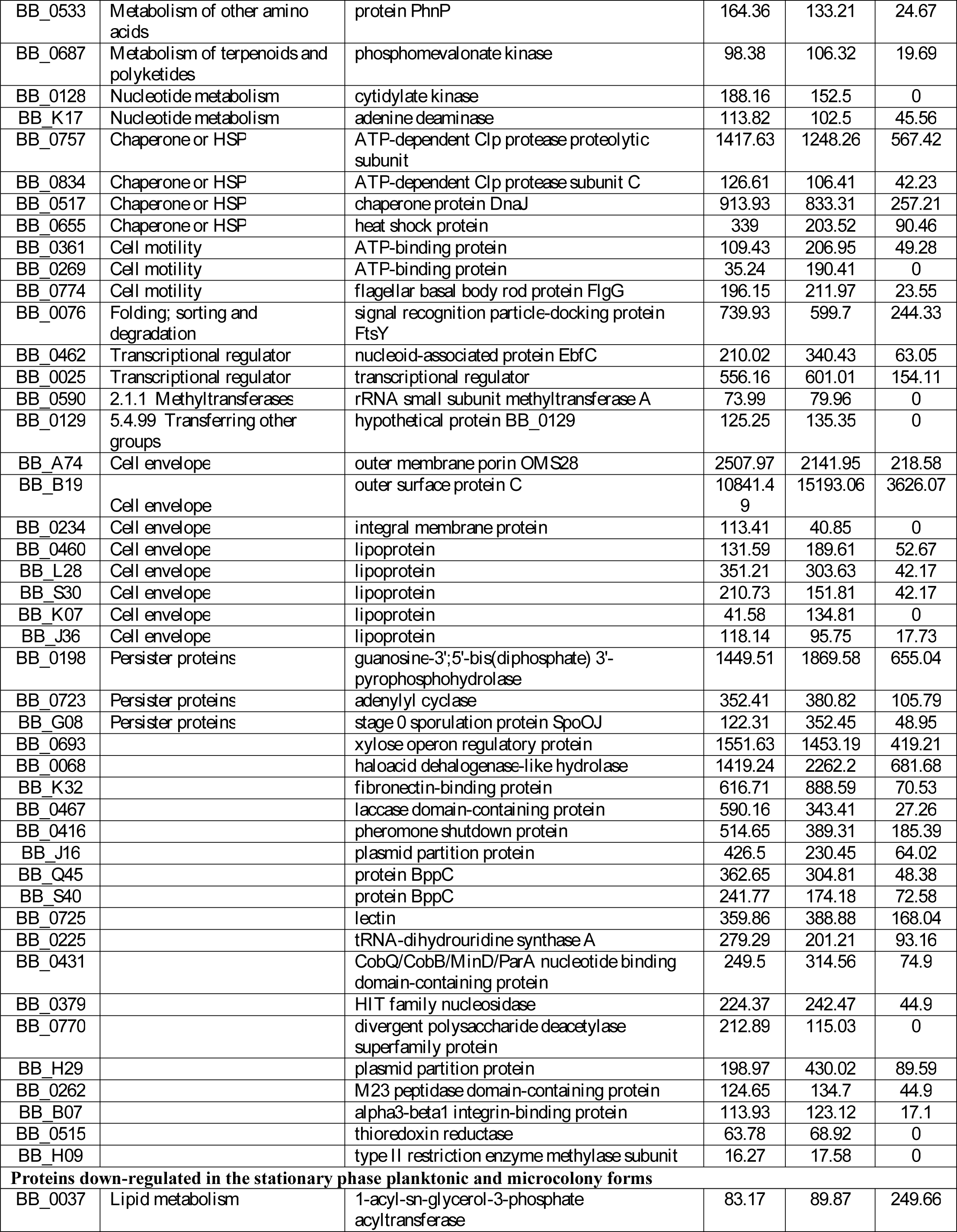
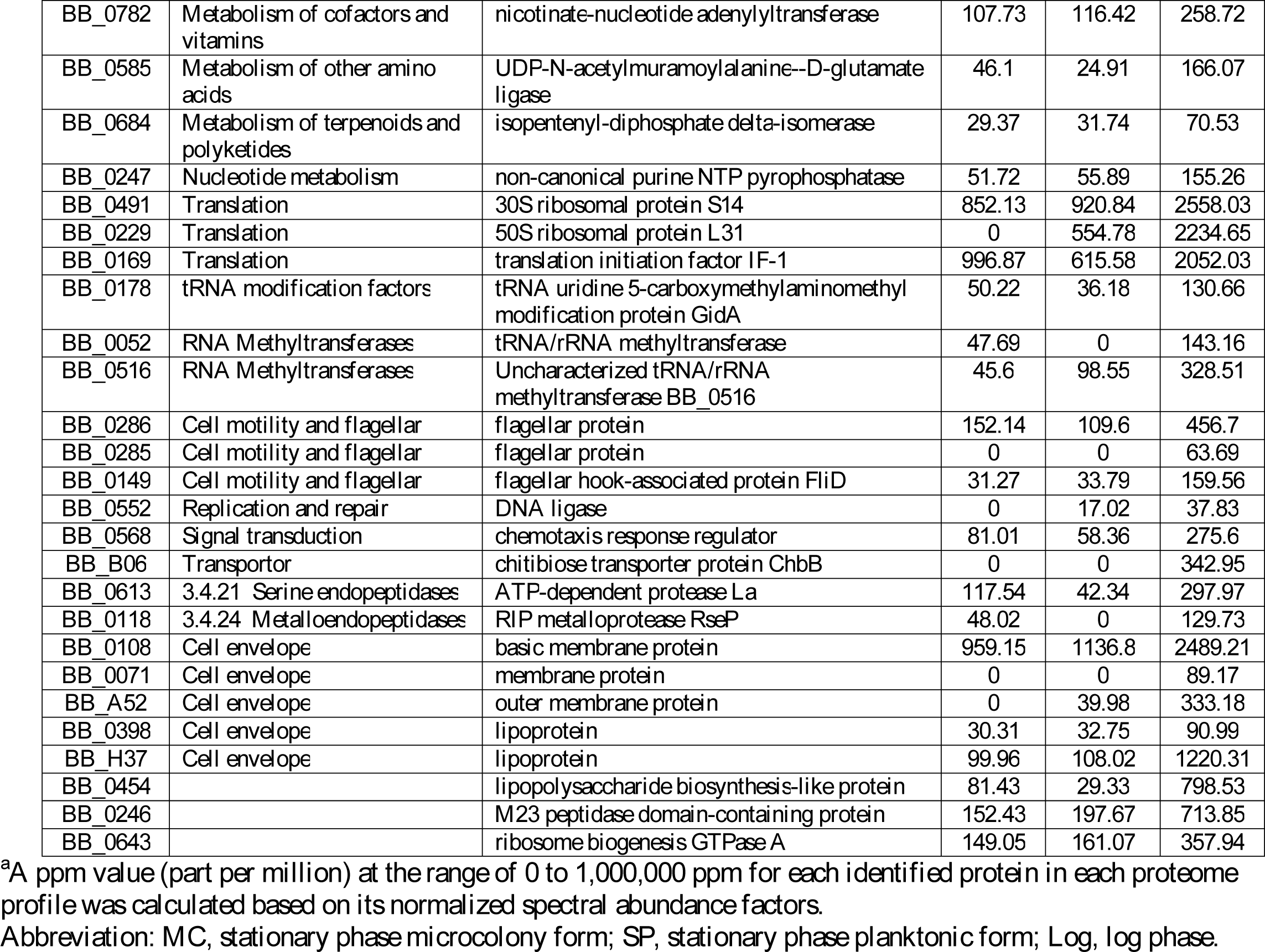
Differentially expressed proteins in both microcolony (MC) form and planktonic spirochete stationary phase (SP) form in a 10-day old culture compared with a 3-day old log phase (Log) spirochete form of *B. burgdorferi*.

**Table 2.**
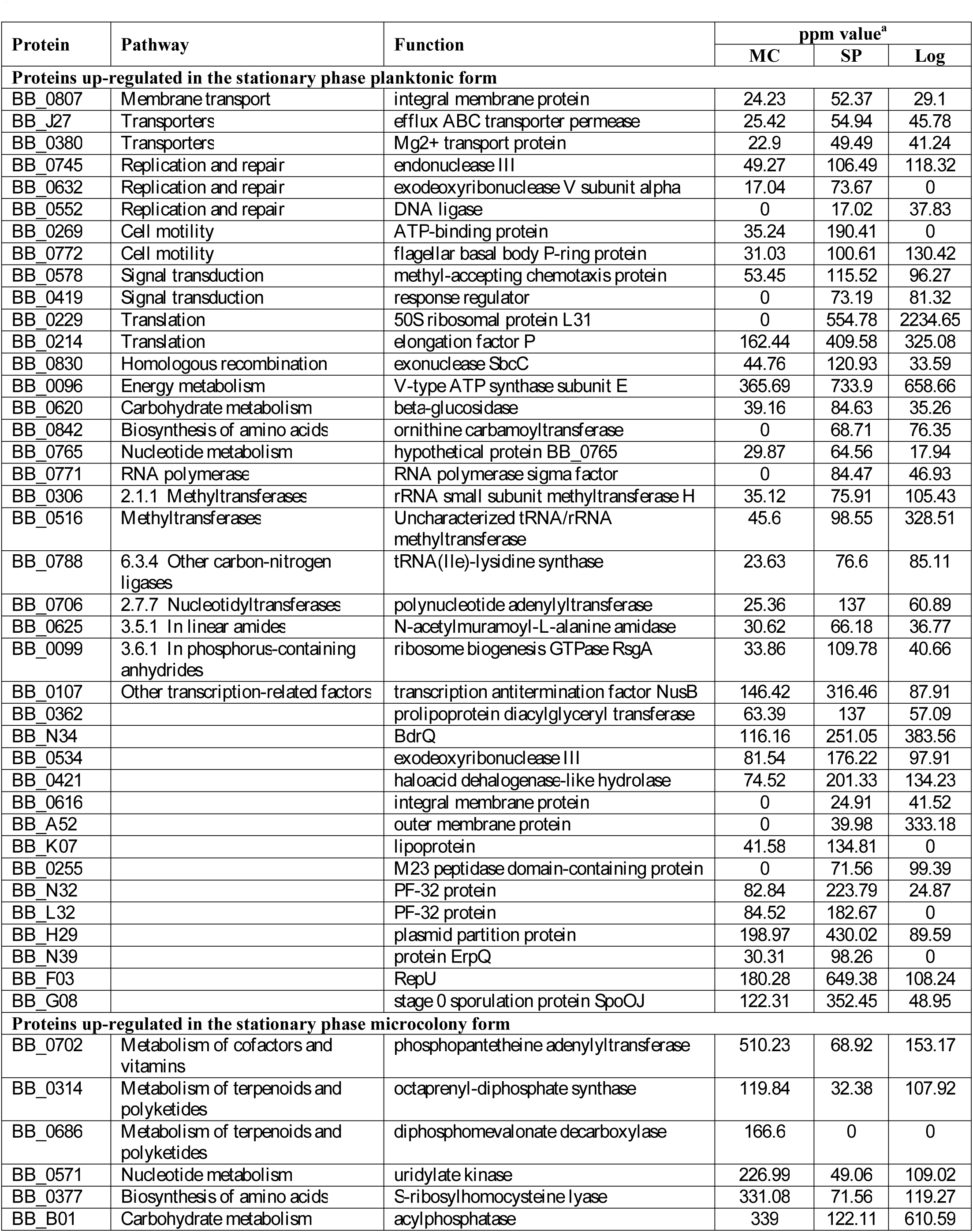
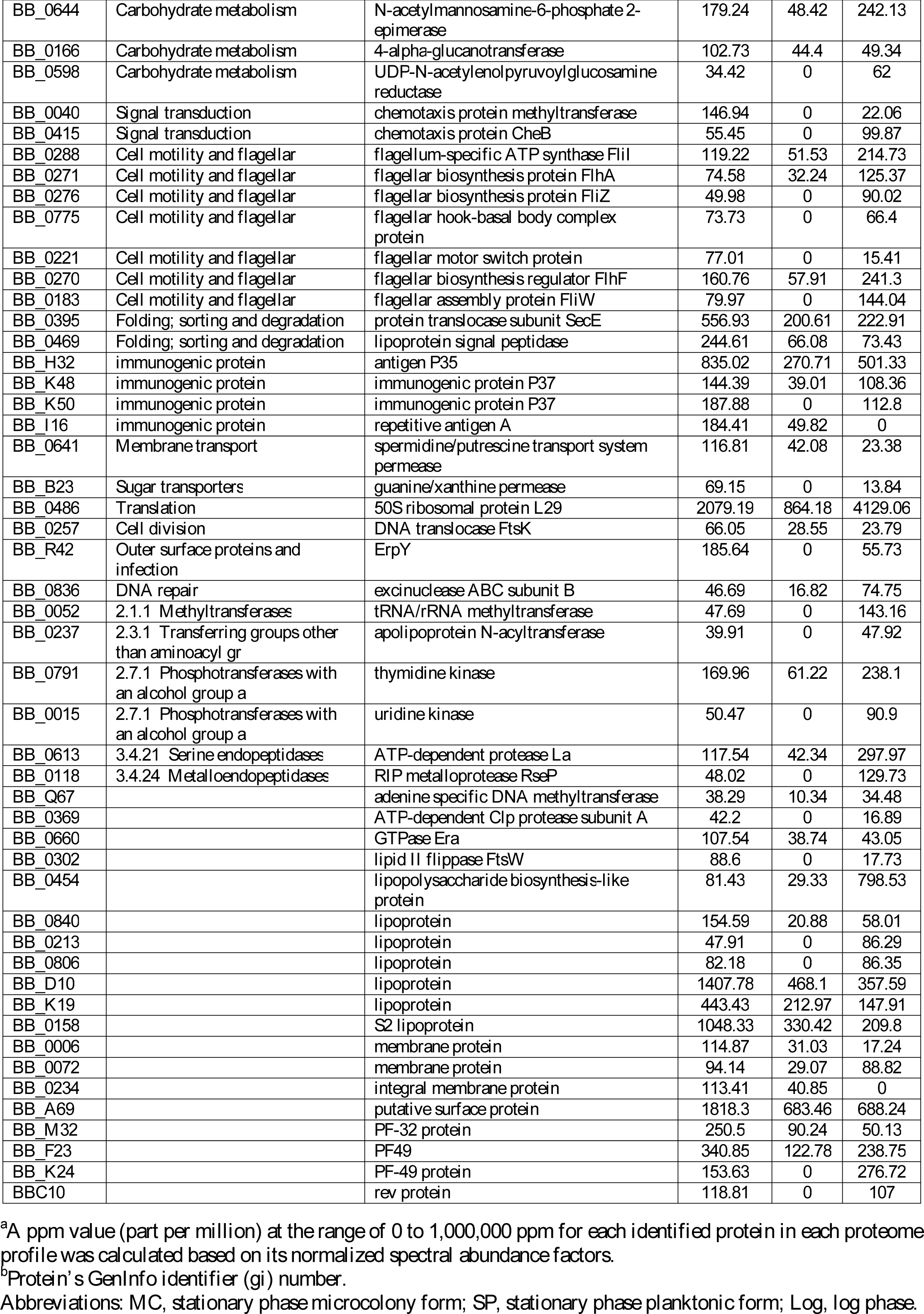
Differentially expressed proteins between the microcolony form and the planktonic form of a 10-day old *B. burgdorferi* stationary phase culture.

### Common altered proteins and pathways shared in stationary phase planktonic form and microcolony form *B. burgdorferi*

Although our previous study identified the gene expression profiles by RNA-seq analysis of *B. burgdorferi* persisters, the proteomic analysis of *B. burgdorferi* persisters has not been performed. To shed new light on the mechanisms of persistence in *B. burgdorferi*, here we compared the proteomic profiles of stationary phase planktonic form and aggregated microcolony cells with log phase spirochetal cells that contain little or no persisters as a control. Compared with the log phase spirochetal cells (>2-fold change, CV>75%, >3 PSMs), a total of 143 proteins were upregulated in both stationary phase forms (MC and SP). Among the 143 core upregulated proteins, 90 proteins had predicted functions and were mapped to different pathways (Table 1, Figure 2). On the other hand, we found that 48 proteins were down-regulated in both MC and SP samples, among which 27 proteins had predicted function and mapped to different pathways (Figure 2). The main up-regulated pathways are involved in infection and virulence, ribosome biogenesis, DNA replication and repair, cell envelope and metabolism. We found 11 proteins associated with infection or virulence (CRASP BB_A68, ErpQ, BdrEFMRVW, DbpAB, BB_I29) are regulated in both stationary phase forms, among them ErpQ protein was only detected in the stationary phase form but not in the log phase. Additionally, 7 up-regulated DNA replication and repair proteins may play an important role in both stationary phase forms for persistence. We also found 6 transporter proteins up-regulated for nutrient transport.

Interestingly, the metabolism of stationary phase *B. burgdorferi* did not seem to be completely dormant in both stationary phase forms with 10 up-regulated and 5 down-regulated metabolism proteins (Table 1, Figure 2). In the two stationary phase forms, the cell envelope structure may be modified because we found 8 cell envelope proteins up-regulated and five down-regulated (Table 1). Besides, we also noticed that 4 molecular chaperone or heat shock proteins including ClpC (BB_0834), ClpP (BB_0757), DnaJ (BB_0517) and heat shock protein BB_0655 were significantly up-regulated in both stationary phase forms. Also, three proteins (BB_0723, BB_0198 SpoT and BB_G08 SpoOJ) up-regulated in both stationary phase forms may play a role in the persistence. Additionally, the up-regulated proteins in both stationary phase forms also include 2 transcriptional regulators (EbfC and BB_0025) and one immunogenic protein BB_I16.

Compared to the previous RNA-seq study on antibiotic tolerant *B. burgdorferi* persisters [15], we can find some common up-regulated pathways between the antibiotic tolerant persisters and the two stationary phase forms (Figure 2). Firstly, some infection and virulence proteins like DbpAB are up-regulated in both antibiotic tolerant and stationary phase *B. burgdorferi* cells. Secondly, it is worth noting that some molecular chaperone or heat shock proteins are up-regulated in these persister forms, such as ClpP and ClpC proteases, hear shock protein DnaJ and GrpE. Besides, DNA repair proteins and some nutrient transporters are also up-regulated in both drug tolerant persisters and stationary phase *B. burgdorferi*.

### Comparison of proteome profile of microcolony form and stationary phase planktonic form of *B. burgdorferi*

Although half (50.1%) of the differentially expressed proteins in stationary phase planktonic form and microcolony form are shared, we found some differences between the proteome profiles of these two forms. For the two stationary phase forms, a total of 146 proteins were differentially expressed (>= 2-fold change, CV>75%, >3 PSMs) between the planktonic form and the microcolony form, among them 58 proteins were up-regulated in the planktonic form and 87 proteins were up-regulated in the microcolony form. There are 39 up-regulated proteins (Table 2) with predicted functions in the stationary phase planktonic form, which includes 3 transporter proteins (BB_0380, BB_J27, BB_0807), 3 DNA replication and repair proteins, 4 RNA polymerase and modification enzymes (BB_0771, BB_0516, BB_0306, BB_0788), 4 motility and signal transduction proteins (BB_0578, BB_0419, BB_0269, BB_0772), 3 cell envelop proteins (BB_0616, BB_A52, BB_K07) and some other proteins of unknown function. On the other hand, 55 up-regulated proteins with predicted functions in the microcolony form mainly include 9 metabolism proteins, 7 flagella proteins (BB_0221, BB_0183, BB_0270, BB_0775, BB_0271, BB_0276, BB_0288), 10 cell envelope proteins including 6 lipoproteins, 4 possible immunogenic proteins (BB_H32, BB_K48, BB_K50, BB_I16), 2 transporter proteins (BB_0641, BB_B23) and 3 DNA/RNA modification proteins (BB_Q67, BB_0836, BB_0052) (Table 2).

## Discussion

Proteomics approach has been used previously to study the *B. burgdorferi* proteome difference during transmission between ticks and mammals [27], in response to environmental variations of temperature and pH [28], and between various native species [29]. However, previous studies provided little quantitative information, and did not compare the proteomes of morphological variants of *B. burgdorferi* in different growth period. In this study, we successfully detected about 80% (1023 proteins) of the 1283 genomically encoded proteins from three forms, i.e. log phase spirochetal form, stationary phase planktonic form, and stationary phase microcolony form of *B. burgdorferi*. This high coverage could display the dynamics of protein expression in different forms of *B. burgdorferi*. Meanwhile, most of selected house-keeping proteins were at similar level or the changes were less than 50% among the three different forms, indicating that the coverage of total proteins identified in each of the three forms is very good and the results are credible.

Compared with the log phase culture, stationary phase planktonic and microcolony forms shared many common features (Figure 2). An important observation of this study is that virulence and infection related proteins are significantly upregulated in the stationary planktonic and microcolony forms compared with log phase spirochete form. In particular, we identified 12 proteins (BB_A07, BB_A68, BB_I29, ErpQ, DbpAB and BdrEFMRVW) up-regulated in the stationary planktonic and microcolony forms involved in infection or virulence of *B. burgdorferi* (Table 1, Figure 2). The category of infection or virulence possesses most up-regulated proteins of both stationary phase forms (Figure 2). And the other four Bdr proteins (BdrAPOH) were up-regulated in the stationary phase planktonic form (Table S1, Figure 2). Protein ChpAI (BB_A07) as a surface lipoprotein plays an important role in the transmission of *B. burgdorferi* from the tick to the mammalian host [30]. Proteins CRASP BB_A68 [31, 32], ErpQ [33], DbpA (BB_A24) and DbpB (BB_A25) [34] could bind to host proteins and play a role in the infection and/or elicit immune response. In our RNA-seq study, *B. burgdorferi* decorin-binding proteins (Dbp) A and B are also up-regulated in the amoxicillin tolerant persisters. The up-regulated expression of these binding proteins in stationary phase cells may strengthen interaction with extracellular proteins and promote aggregation and resistance to stress environment. Protein BB_I29, a virulence associated lipoprotein, also up regulated 6-8 fold in the two stationary phase from. Additionally, we found a fibronectin-binding protein BB_K32 was up regulated about 10 fold in the stationary phase forms compared to log phase form (Table 1). Protein BB_K32 as an antigen elicited protective immunity in mice was detected in spirochetes infecting mice but not in spirochetes grown *in vitro* [35, 36]. These results showed that the stationary phase cells *in vitro* might have some common features with the *in vivo* infecting cells and suggest that stationary phase *B. burgdorferi* culture would be a better *in vitro* model for persistence and virulence studies and drug screens [9] than the log phase cells. Indeed, this is consistent with our recent study which shows that stationary phase and microcolony forms of *B. burgdorferi* produce more severe lesions and arthritis than log phase spirochetal form in the mouse model (https://www.biorxiv.org/content/early/2018/11/28/440461).

Additionally, consistent with the findings of our previous drug tolerant *B. burgdorferi* persister RNA-seq study [15], we found some DNA repair related proteins (BB_0632, BB_0111, BB_0114, BB_0797, BB_0623, and BB_0837) were also up-regulated in the two stationary phase forms, which would help maintain stability of DNA under starvation and stress conditions (Figure 2). Furthermore, we found that six membrane transporter proteins (BB_0604, BB_0217, BB_0219, BB_0334, BB_0574, and BB_A34) involved in phosphate transport system, peptide uptake transporter, extracellular solute transporter and metal cation transporter were up-regulated in the stationary phase cells. These transporters could uptake nutrients and regulate intracellular ion concentration to cope with the starvation environment and could be important for in vivo survival and virulence.

In this study, we found four heat shock proteins (ClpC BB_0834, ClpP BB_0757, DnaJ BB_0517, BB_0655) including two Clp protease proteins which were up-regulated in both stationary phase forms (Table 1, Figure 2). We also noticed that *clp* gene which encodes protease necessary for degrading toxic proteins was one of the most highly upregulated gene in doxycycline and amoxicillin treated *B. burgdorferi* persisters in our previous RNA-seq study [15]. These up-regulated heat shock proteins or protein degradation pathway proteins could stabilize new proteins or detoxify toxic protein build-up in persister cells to ensure correct folding and help to refold or degrade damaged proteins under stress, which could be an important mechanism for persister survival. This is consistent with the previous observations that protein degradation pathways such as trans-translation and Clp proteases are critical for persister survival (Shi et al., 2011; Zhang S 2017).

In addition, we found the cell envelope proteins changed greatly in both stationary phase forms compared to log phase cells; 8 envelope proteins were up-regulated, and 5 proteins were down-regulated in both stationary phase samples. Up-regulated cell structure proteins include five lipoproteins, three bacterial envelope proteins including outer surface protein C (OspC, Table 1). OspC plays an important role in the *B. burgdorferi* transmission from tick to mammalian host [32]. And OspC as a virulence factor is essential for infection in the mammalian host, as OspC deletion mutant was not able to infect naive mice [37]. The down-regulated cell structure proteins include three lipoproteins, three bacterial membrane proteins, and three motility proteins. For the round body form in the stationary phase cells, changing component of membrane proteins could stabilize inner membrane structure to make up defect of outer membrane. Also, the aggregated microcolony could change the structure of cell envelope to adapt to the stress condition. Changes in cell envelope would be one of the causes for stationary phase cell tolerance to antibiotics.

Compared to the log phase culture which contains primarily spirochete form, stationary phase *B. burgdorferi* culture consists of multiple morphological forms including planktonic form (spirochetes and round-body) and aggregated microcolony form [9]. In this study, we separated these two forms (planktonic and microcolony) from the stationary phase culture and compared their protein expression. Despite the morphological structure of these two forms is very different (Figure 1B and 1C), their proteome profile was changed only slightly. Only 19.4% proteins (146 in 750 proteins) were differentially expressed (>= 2-fold change, CV>75%, >3 PSMs) between the stationary phase planktonic form and microcolony form. Analysis of 94 differentially expressed proteins showed that the proteome difference is mainly in the cell envelope proteins, flagella proteins and metabolism (Figure 2). Compared to the log phase and stationary phase planktonic form, this study found 9 cell envelope proteins and 5 signal transduction proteins (two chemotaxis CheY proteins, chemotaxis methyltransferase, methyl-accepting chemotaxis protein and purine-binding chemotaxis protein) that were up regulated only in the aggregated microcolony form (Table S2, Figure 2). This suggests the aggregated microcolony form of *B. burgdorferi* possesses specific cell envelope structure and has more contact with surrounding environment and adjacent cells, compared with the log phase and stationary phase planktonic form. Among the different morphological forms, aggregated microcolony form is the most drug tolerant form [12]. The drug tolerance of the microcolony form may be related to the differences in specific cell envelope structure and cell contact. We identified 10 uniquely down-regulated flagella proteins in the stationary phase planktonic form of *B. burgdorferi* compared with log phase (Table S1, Figure 2). Flagella proteins of *B. burgdorferi* are not only responsible for the cell motility but also the cell structure [38]. Stationary phase planktonic form of *B. burgdorferi* down-regulated these flagella proteins which may change their cell structure and morphology and may explain the formation of variant forms like round-bodies and microcolonies in the *B. burgdorferi* stationary phase [9, 12]. However, interestingly, we still observed some flagella proteins being upregulated in the microcolony form compared with planktonic stationary phase *B. burgdorferi* (Table 2). This is consistent with our finding that some active spirochetes are located on the surface of the aggregated microcolony form (data not shown).

Remarkably, three persister proteins (BB_0723 adenylyl cyclase, SpoT guanosine-3’;5’-bis(diphosphate) 3’-pyrophosphohydrolase and sporulation protein spoOJ) were up-regulated in both stationary phase forms (Figure 2, Table 1), and another two persister related GTPases Era and Obg were found to be up-regulated in the microcolony form (Figure 2, Table S2). Adenylyl cyclase can synthesize cAMP which is involved in the formation of bacterial persisters [20]. Enzyme SpoT plays a role in the guanosine tetraphosphate (ppGpp) mediated persistence [24–26]. In addition, two GTPases Era [21, 22] and Obg [23] up-regulated in the microcolony form may also play an important role in the bacterial persistence. These findings provide some explanation of why the stationary phase variant forms especially the microcolony form was more persistent than the log phase growing spirochetal form [12, 39].

Bacterial cellular metabolism is known to affect persister levels [40, 41]. It is of interest to note that in the two stationary phase forms, we found 10 metabolism proteins were up-regulated compared to the log phase cells (Table 1, Figure 2, Table S1). Among them, four significantly up-regulated (>5 fold) proteins are associated with carbohydrate metabolism (BB_0364, methylglyoxal synthase), energy (BB_0092, V-type ATPase subunit D), other amino acid (BB_0533, PhnP protein), and nucleotide (BB_0128, cytidylate kinase) (Table 1). Interestingly, we noticed methylglyoxal synthase [42], PhnP protein [43] and cytidylate kinase [44] are all involved in phosphoryl transfer and phosphate metabolism. These suggest phosphate metabolism may also play a role in the *B. burgdorferi* persister formation as in *E. coli* [40]. In addition, one arginine metabolism protein, ornithine carbamoyltransferase (BB_0842), was significantly down-regulated (<5 fold), which is consistent with previous studies demonstrating that arginine metabolic pathway may help persisters survive under stress conditions [45, 46](https://www.biorxiv.org/content/early/2017/03/07/114827). Not all the metabolic genes are down-regulated in the persisters although the metabolism level of the dormant form are lower than the growing form. Meanwhile, we also found while some ribosomal proteins were down-regulated, some others were up-regulated in both stationary phase forms (Figure 2, Table 1). In a recent study, *rpmF* encoding a ribosomal protein was also identified as a persister gene in *E. coli* [47], which supports the possibility that certain ribosomal proteins may play a role in persister formation or survival in *Borrelia* SP and MC forms. In future studies we will confirm whether these proteins or genes are involved in persistence to antibiotics and multiple stresses.

This study not only uncovered pathways by which *B. burgdorferi* copes with stress and morphs into stress-resistant forms but also shed new light on mechanisms of persistence in this versatile pathogen with implications for understanding disease pathogenesis. The proteome profile of the stationary phase planktonic and microcolony forms showed some common features with the *in vivo* infecting bacteria since infection and virulence related proteins are upregulated in these variant forms (Figure 2). This finding supports that stationary phase *B. burgdorferi* may offer a reasonable *in vitro* model for *in vivo* drug screens. In the meantime, some proteins and pathways identified in this study need further studies such as overexpression or knockout to verify their roles in persistence and virulence. The confirmed crucial proteins by *in vitro* and *in vivo* studies could be good candidates for drug development for improve treatment of Lyme disease. Moreover, the proteins identified to be preferentially overexpressed in variant persister forms such as round bodies and microcolonies could serve as antigens for vaccines and for improved diagnosis of Lyme disease. Indeed, our recent study has shown that inclusion of antigens from these variant forms could improve the sensitivity of the current ELISA test in diagnosis of Lyme disease (M Cai et al. and Y Zhang to be published).

## Acknowledgments

We acknowledge the support by Global Lyme Alliance.

## References

1. CDC. Lyme Disease. 2017 7/3/2018]; Available from: http://www.cdc.gov/lyme/.

2. CDC. Post-Treatment Lyme Disease Syndrome. 2017 7/3/2018]; Available from: http://www.cdc.gov/lyme/postLDS/index.html.

3. Hodzic, E., et al., Resurgence of Persisting Non-Cultivable Borrelia burgdorferi following Antibiotic Treatment in Mice. PLoS One, 2014. 9(1): p. e86907.

4. Embers, M.E., et al., Persistence of Borrelia burgdorferi in Rhesus Macaques following Antibiotic Treatment of Disseminated Infection. PLoS One, 2012. 7(1): p. e29914.

5. Hodzic, E., et al., Persistence of Borrelia burgdorferi following antibiotic treatment in mice. Antimicrob Agents Chemother, 2008. 52(5): p. 1728–36.

6. Straubinger, R.K., et al., Persistence of Borrelia burgdorferi in experimentally infected dogs after antibiotic treatment. Journal of clinical microbiology, 1997. 35(1): p. 111–6.

7. Marques, A., et al., Xenodiagnosis to Detect Borrelia burgdorferi Infection: A First-in-Human Study. Clin Infect Dis, 2014. 58(7): p. 937–45.

8. Crossland, N.A., X. Alvarez, and M.E. Embers, Late Disseminated Lyme Disease: Associated Pathology and Spirochete Persistence Posttreatment in Rhesus Macaques. Am J Pathol, 2018. 188(3): p. 672–682.

9. Feng, J., et al., Identification of novel activity against Borrelia burgdorferi persisters using an FDA approved drug library. Emerg Microbes Infect, 2014. 3(7): p. e49.

10. Sharma, B., et al., Borrelia burgdorferi, the Causative Agent of Lyme Disease, Forms Drug-Tolerant Persister Cells. Antimicrob Agents Chemother, 2015. 59(8): p. 4616–24.

11. Caskey, J.R. and M.E. Embers, Persister Development by Borrelia burgdorferi Populations In Vitro. Antimicrob Agents Chemother, 2015. 59(10): p. 6288–95.

12. Feng, J., P.G. Auwaerter, and Y. Zhang, Drug Combinations against Borrelia burgdorferi persisters In Vitro: Eradication Achieved by Using Daptomycin, Cefoperazone and Doxycycline. PLoS One, 2015. 10(3): p. e0117207.

13. Miklossy, J., et al., Persisting atypical and cystic forms of Borrelia burgdorferi and local inflammation in Lyme neuroborreliosis. J Neuroinflammation, 2008. 5: p. 40.

14. Lantos, P.M., P.G. Auwaerter, and G.P. Wormser, A systematic review of Borrelia burgdorferi morphologic variants does not support a role in chronic Lyme disease. Clin Infect Dis, 2014. 58(5): p. 663–71.

15. Feng, J., et al., Persister mechanisms in Borrelia burgdorferi: implications for improved intervention. Emerg Microbes Infect, 2015. 4(8): p. e51.

16. Florens, L., et al., Analyzing chromatin remodeling complexes using shotgun proteomics and normalized spectral abundance factors. Methods, 2006. 40(4): p. 303–311.

17. Paoletti, A.C., et al., Quantitative proteomic analysis of distinct mammalian Mediator complexes using normalized spectral abundance factors. Proceedings of the National Academy of Sciences of the United States of America, 2006. 103(50): p. 18928–18933.

18. Kanehisa, M., et al., KEGG: new perspectives on genomes, pathways, diseases and drugs. Nucleic Acids Res, 2017. 45(D1): p. D353-D361.

19. Altschul, S.F., et al., Gapped BLAST and PSI-BLAST: a new generation of protein database search programs. Nucleic Acids Res, 1997. 25(17): p. 3389–402.

20. Amato, S.M., M.A. Orman, and M.P. Brynildsen, Metabolic control of persister formation in Escherichia coli. Mol Cell, 2013. 50(4): p. 475–87.

21. Auvray, F., et al., The Listeria monocytogenes homolog of the Escherichia coli era gene is involved in adhesion to inert surfaces. Appl Environ Microbiol, 2007. 73(23): p. 7789–92.

22. Ji, X., Structural insights into cell cycle control by essential GTPase Era. Postepy Biochem, 2016. 62(3): p. 335–342.

23. Verstraeten, N., et al., Obg and Membrane Depolarization Are Part of a Microbial Bet-Hedging Strategy that Leads to Antibiotic Tolerance. Mol Cell, 2015. 59(1): p. 9–21.

24. Germain, E., et al., Stochastic induction of persister cells by HipA through (p)ppGpp-mediated activation of mRNA endonucleases. Proceedings of the National Academy of Sciences, 2015. 112(16): p. 5171–5176.

25. Hauryliuk, V., et al., Recent functional insights into the role of (p)ppGpp in bacterial physiology. Nat Rev Microbiol, 2015. 13(5): p. 298–309.

26. Liu, S., et al., Variable Persister Gene Interactions with (p)ppGpp for Persister Formation in Escherichia coli. Front Microbiol, 2017. 8: p. 1795.

27. Norris, S.J., The dynamic proteome of Lyme disease Borrelia. Genome Biol, 2006. 7(3): p. 209.

28. Angel, T.E., et al., Proteome analysis of Borrelia burgdorferi response to environmental change. PLoS One, 2010. 5(11): p. e13800.

29. Schnell, G., et al., Proteomic analysis of three Borrelia burgdorferi sensu lato native species and disseminating clones: relevance for Lyme vaccine design. Proteomics, 2015. 15(7): p. 1280–90.

30. Xu, H., et al., Role of the surface lipoprotein BBA07 in the enzootic cycle of Borrelia burgdorferi. Infect Immun, 2010. 78(7): p. 2910–8.

31. McDowell, J.V., et al., Putative coiled-coil structural elements of the BBA68 protein of Lyme disease spirochetes are required for formation of its factor H binding site. J Bacteriol, 2005. 187(4): p. 1317–23.

32. Kenedy, M.R., T.R. Lenhart, and D.R. Akins, The role of Borrelia burgdorferi outer surface proteins. FEMS Immunol Med Microbiol, 2012. 66(1): p. 1–19.

33. Miller, J.C., et al., Temporal analysis of Borrelia burgdorferi Erp protein expression throughout the mammal-tick infectious cycle. Infect Immun, 2003. 71(12): p. 6943–52.

34. Salo, J., et al., Decorin binding by DbpA and B of Borrelia garinii, Borrelia afzelii, and Borrelia burgdorferi sensu Stricto. J Infect Dis, 2011. 204(1): p. 65–73.

35. Fikrig, E., et al., Borrelia burgdorferi P35 and P37 proteins, expressed in vivo, elicit protective immunity. Immunity, 1997. 6(5): p. 531–9.

36. Suk, K., et al., Borrelia burgdorferi genes selectively expressed in the infected host. Proc Natl Acad Sci U S A, 1995. 92(10): p. 4269–73.

37. Tilly, K., et al., Borrelia burgdorferi OspC protein required exclusively in a crucial early stage of mammalian infection. Infect Immun, 2006. 74(6): p. 3554–64.

38. Motaleb, M.A., et al., Borrelia burgdorferi periplasmic flagella have both skeletal and motility functions. Proc Natl Acad Sci U S A, 2000. 97(20): p. 10899–904.

39. Lewis, K., Persister cells, dormancy and infectious disease. Nat Rev Microbiol, 2007. 5(1): p. 48–56.

40. Li, Y. and Y. Zhang, PhoU is a persistence switch involved in persister formation and tolerance to multiple antibiotics and stresses in Escherichia coli. Antimicrob Agents Chemother, 2007. 51(6): p. 2092–9.

41. Amato, S., et al., The role of metabolism in bacterial persistence. Frontiers in Microbiology, 2014. 5(70).

42. Totemeyer, S., et al., From famine to feast: the role of methylglyoxal production in Escherichia coli. Mol Microbiol, 1998. 27(3): p. 553–62.

43. He, S.M., et al., Structure and mechanism of PhnP, a phosphodiesterase of the carbon-phosphorus lyase pathway. Biochemistry, 2011. 50(40): p. 8603–15.

44. Verma, N.K. and B. Singh, Insight from the structural molecular model of cytidylate kinase from Mycobacterium tuberculosis. Bioinformation, 2013. 9(13): p. 680–4.

45. Yee, R., et al., Genetic Screen Reveals the Role of Purine Metabolism in Staphylococcus aureus Persistence to Rifampicin. Antibiotics, 2015. 4(4): p. 627.

46. Wu, N., et al., Ranking of persister genes in the same Escherichia coli genetic background demonstrates varying importance of individual persister genes in tolerance to different antibiotics. Frontiers in Microbiology, 2015. 6(1003).

47. Liu, S., et al., Identification of novel genes including rpmF and yjjQ critical for Type II persister formation in Escherichia coli. bioRxiv, 2018.

